# RNA-guided nucleases enable a gene drive of insertion sequences in plasmids

**DOI:** 10.1101/2025.02.20.638934

**Authors:** Kepler S. Mears, Fernando W. Rossine, Natalia Quinones-Olvera, Célia Souque, Sophia S. Wiesenfeld, Kesther D. C. Jean, Shreyas V. Pai, Michael Baym

**Affiliations:** Department of Biomedical Informatics and Laboratory of Systems Pharmacology, Harvard Medical School, Boston, MA, USA; Roxbury Community College, Boston, MA, USA

**Author notes:** These authors contributed equally (Kepler S. Mears, Fernando W. Rossine).

## Abstract

Mobile genetic elements (MGEs) are diverse, self-replicating DNA molecules that can reside within cellular hosts and integrate into one another. This co-occurrence imposes distinct evolutionary pressures. Plasmids often contain insertion sequences (ISs). However, the multi-copy nature of plasmids should hinder IS introduction and spread, disfavoring inheritance of nascent plasmid variants through genetic drift. Mechanisms by which ISs overcome these barriers remain unidentified. Here we find that the RNA-guided nuclease TnpB enables such a mechanism to bias its inheritance in plasmids. We show that TnpB, the likely ancestor to Cas12, enables a gene drive to spread the IS within multicopy plasmids and functions as a primitive anti-self defense system in conjugative plasmids. The gene drive between TnpB-bearing ISs and plasmids promotes the spread of both MGEs beyond the ability of either individually. The nested existence between MGEs is not an incidental result of selfish spread, but a driver of it.

## Introduction

Mobile genetic elements (MGEs) are a diverse group of DNA molecules that can replicate independently of the host chromosome, and therefore face evolutionary pressures of their own ^1,2^. For example, plasmids are circular, extrachromosomal, self-replicating DNA molecules, often capable of engaging in horizontal gene transfer (HGT) between hosts ^3^. In contrast, transposons (TNs) are linearly integrated segments of DNA that can jump between genomic contexts and rarely engage in HGT directly ^4,5^. Notably, MGEs are a major source of evolutionary novelty in bacteria: plasmids can transfer large amounts of genetic material between bacterial lineages while TNs can disrupt genes upon insertion and modulate transcription. Despite their different structures and functions, MGEs are often found in nested configurations. Plasmids can carry TNs and are particularly enriched in insertion sequences (ISs) (**Figure 1A**), minimal TNs carrying only the necessary genes for their transposition and maintenance ^4–7^. The intimate relationship between ISs and plasmids raises the question of what evolutionary processes lead to their association and whether this relationship drives bacterial genome plasticity beyond the abilities of either MGE individually.

**Figure 1.**
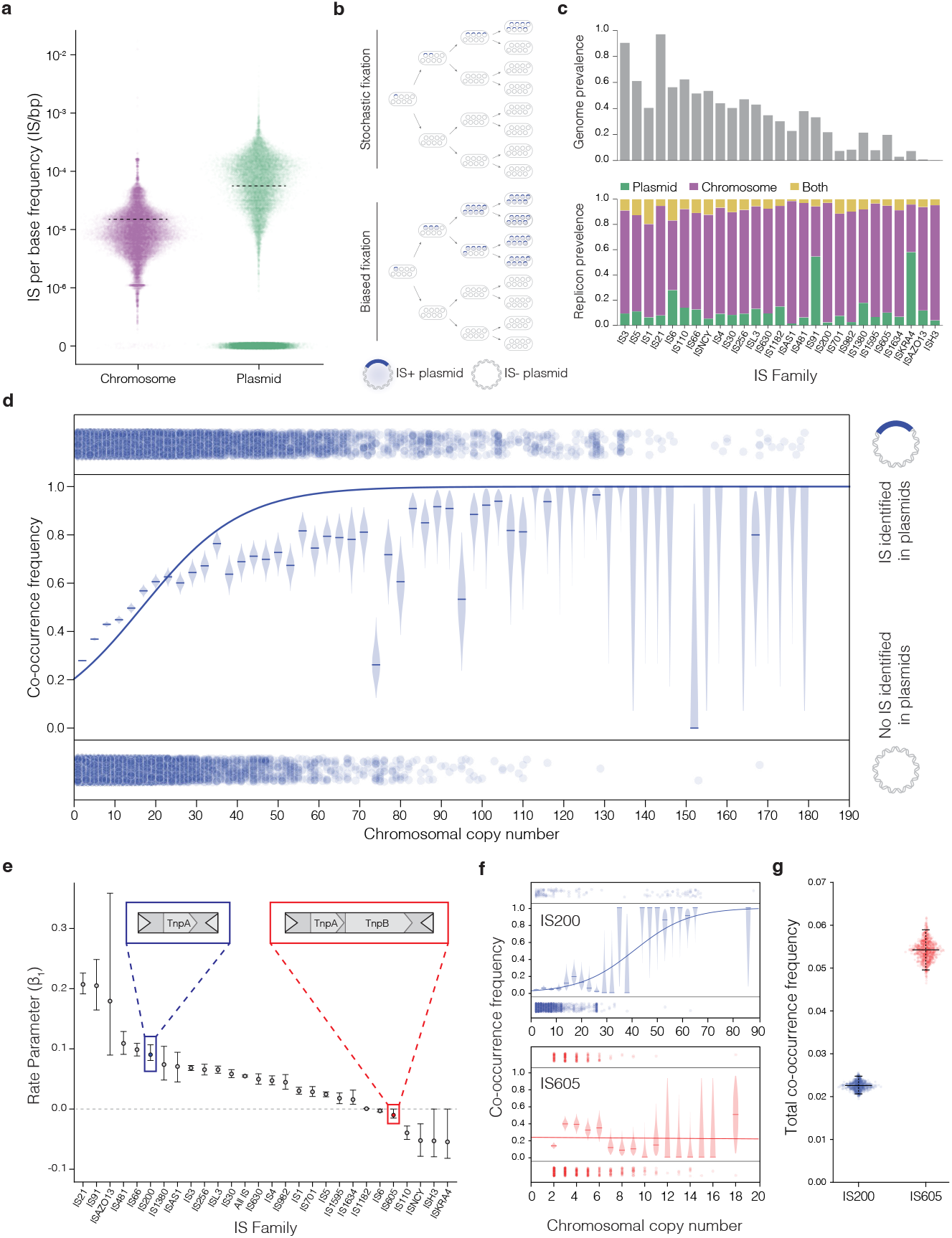
Positive association between chromosomal copy number and plasmid co-occurrence is a potential general mechanism to overcome limits in stochastic fixation for insertion sequences. A) Per base frequency of insertion sequences in plasmids and chromosomes in the PLSDB database, the dotted line indicates the median. B) Model of purely stochastic IS fixation in plasmids vs that of an IS with increased fixation rate. IS+ plasmids are indicated in blue and IS- in grey C) Prevalence of insertion sequences across genomes in the PLSDB database (top) and distribution across replicon types (bottom). Insertion sequence families are ordered from highest overall abundance to lowest (Supplemental Figure 1B). D) Co-occurrence frequency of insertion sequence in plasmids and chromosomes vs IS chromosomal copy numbers. Dots on the bottom represent an IS found only in the chromosome of a genome while dots on top represent IS that co-occur in the chromosome and plasmid. Binned and calculated frequencies at specified chromosomal copy number are plotted with the expected beta-posterior beta distribution in light blue and medians indicated with a horizontal line. The sigmoid fit determined from the raw data is plotted overtop in solid blue. E) Rate parameters from the sigmoid fit of all IS families analyzed with 95% confidence intervals determined from a chi-squared approximation from a profile likelihood of the rate parameter. F) Individual co-occurrence frequency plots with overlaid sigmoid fit for IS200 in blue, IS605 in red, with corresponding binned frequencies, expected posterior and fit. G) Overall co-occurrence frequency of plasmid presence for TnpA and TnpA/B IS. 95% confidence intervals determined from bootstrapping (N=1000) with each replicate in light blue/red.

The integration of ISs into plasmids enhances the evolvability of both MGEs and their bacterial hosts. ISs enable plasmids to acquire genes by providing recombination homology regions, diversifying the plasmid gene repertoire ^8–10^. Conversely, plasmids allow ISs to mobilize into different hosts and spread into new genomic contexts ^11^. Altogether, these MGE activities open new evolutionary paths for their bacterial hosts, regardless of any direct fitness effects. However, this increased evolvability is a second-order selective benefit which may not, by itself, explain the spread of IS-bearing (IS+) plasmids ^12,13^. Importantly, theory predicts that genes that increase the evolvability of an organism are unlikely to sweep through a population if they are initially present at a low density ^14^. Therefore, the origin of the high prevalence of IS on plasmids remains unexplained.

IS+ plasmids face distinct challenges spanning many scales of biological organization as they spread across a bacterial population. First, an IS must transpose into an IS-plasmid, which is stoichiometrically disfavored by the higher proportion of chromosomal DNA of most bacterial cells. Second, because plasmids often exist in multiple copies per cell, the nascent IS+ plasmid is in direct intracellular competition with the IS-copies of the plasmid (**Figure 1B, top**). Moreover, recent work suggests that the acquisition of an IS might impair the within-cell replicative fitness of the plasmid ^15^. This decreased replication across cell cycles reduces the probability of a newly divided cell inheriting only IS+ plasmids (i.e. fixing the IS+ plasmid). Finally, any cells that may have fixed the IS+ plasmid must compete against most of the bacterial population that carries only the ancestral IS-plasmid. This competition between cells may further disfavor the spread of the IS+ plasmids, due to a general lack of a host fitness benefit conferred by the IS acquisition. However, these theoretical challenges to the spread of passenger ISs are in stark contrast with the high IS prevalence observed in plasmids.

The discrepancy between the high IS+ plasmid prevalence and the obstacles to their spread can be reconciled since ISs are not passive plasmid passengers. TNs and ISs often encode systems known to improve their chromosomal retention ^16,17^, which could also promote their stability in plasmids. For instance, widely distributed IS families carry TnpBs, which are programmable, RNA-guided endonucleases. These enzymes can cleave DNA sequences that lack the IS but that are homologous to where the IS has inserted. In cases where the TnpB nuclease activity was targeted towards plasmids, TnpB-based systems could favor the fixation probability of IS+ plasmids (**Figure 1B, bottom**). Generally, many such inheritance-biasing mechanisms exist in nature and are a fundamental part of gene drives. Yet, to date, no such gene drive activity, TnpB-dependent or otherwise, has been described for IS+ plasmids.

Here we find both a potential general mechanism to increase IS acquisition by plasmids, and a specific mechanism for ISs to infiltrate plasmid populations. We find a strong association between the chromosomal copy number of an IS and residence on plasmids, suggesting that most IS families rely on high copy numbers to invade plasmids. However, we identify some families that break the trend and appear to possess mechanisms which allow them other means of spread. In particular we find that the IS605 family has a specific mechanism enabled by the TnpB’s RNA-guided nuclease that biases plasmid invasion. The IS605 family utilizes RNA-guided nucleases to enable an intracellular gene drive in multicopy plasmids, promoting the spread of the IS within the plasmid population. This mechanism additionally enables protection of plasmids from other external invaders competing for hosts. The unique association between IS605 and plasmids is the first known instance of a specific mechanism that overcomes the obstacles of plasmid residency by transposons.

## Results

### Insertion sequence chromosomal copy number is predictive of plasmid carriage

Insertion sequences can exist in multiple copies within a bacterial genome. The number of identical copies of an IS in a chromosome, the chromosomal copy number (CCN), is highly flexible and can change as the IS transposes or is lost. More copies of an IS should theoretically result in more potential transposition events, effectively modulating the IS activity which can be further regulated by specific IS mechanisms ^4,5^. We hypothesized that greater IS activity (i.e. greater CCN) would allow for more transposition events from the chromosome into plasmids and therefore result in a greater likelihood of IS co-occurrence (presence of the IS on a plasmid and the chromosome)

To determine IS co-occurrence we searched 19,658 complete contiguous genomes from the PLSDB database of plasmids for insertion sequences and determined their replicon location (chromosome or plasmid) ^18,19^ (**Supplemental Figure 1A**). ISs were very common in plasmids. Out of the total 59,895 plasmids within the dataset, 32,641 contained or more IS elements. Plasmids were also more enriched in ISs having roughly an order of magnitude greater per base frequency compared to the chromosome (**Figure 1A**). We found IS3 to be the most abundant IS with over 250,000 instances in the dataset, (**Supplemental Figure 1B**) while IS21 was the most commonly occurring in genomes with over 90% of all genomes containing one or more instances (**Figure 1C, top**). The majority of IS elements were identified in the chromosome, aside from IS91 and ISKRA4 which were predominantly located on plasmids (**Figure 1C, bottom**).

IS co-occurrence is positively correlated with IS chromosomal copy number (**Figure 1D**). For ISs within a genome we determined the chromosomal copy number (number of identical copies of an IS) and the replicon location (chromosome or plasmid). We then asked whether that same IS also occurred in any of the plasmids within the genome and classified these events as IS co-occurrence (**Figure 1D**, dots bottom and top respectively). We calculated the co-occurrence frequencies at binned chromosomal copy numbers and observed a clear positive association which was reasonably modeled by a logistic regressor (**Figure 1D**).

The majority of IS families followed a similar trend of positive association between chromosomal copy number and IS co-occurrence. To summarize the association across IS families we extracted the rate parameter, β_1_, derived from the sigmoid function with β_1_<0 indicating negative association with chromosomal copy number and β_1_>0 a positive association. As expected, (**Figure 1D**), the majority of IS families had positive rate parameters **(Figure 1E**) which could be readily observed in their individual sigmoid fits, (**Supplemental Figure 1C**) with a handful of outliers having β_1_<0 or β_1_≈0. A few outliers, notably ISH3 and ISKRA4, were likely due to the minimal data and abundance of these IS preventing an association being derived with confidence (**Supplemental Figure 1C**). Increasing chromosomal copy number, and thus IS activity, is a potentially common method for IS to bias their enrichment in plasmids.

### TnpB breaks the positive association between plasmid co-occurrence and IS chromosomal copy number

Among the associations in different IS families we noticed a difference between two highly related families IS200 and IS605 ^20,21^. The IS200 family are some of the most minimal insertion sequences containing only a transposase TnpA while the IS605 (and the similar IS607) family differ by the presence of TnpB, a second gene. The IS200 subfamily followed the general trend with β_1_>0 while IS605 did not with a β_1_≈0. The difference was readily apparent in the respective sigmoid fits for each with IS (**Figure 1F**). Specifically at low chromosomal copy number IS605 had much higher co-occurrence frequency than IS200 implying that IS605 elements have increased fixation rate at low chromosomal copy number compared to IS200 (**Supplemental Figure 1D**). As IS200 and IS605 utilize the same transposition mechanism, we can infer that any difference in interactions between IS and plasmids is likely due to the activity of TnpB.

The mechanism of TnpB has recently been described to aid in genomic retention of the IS ^22^. The TnpA of IS200/605 elements typically target specific, short (4-6 nucleotide), A/T rich motifs and transposes via a scarless cut-and-paste mechanism triggered by genome replication ^20^. The result of excision leaves one daughter chromosome without the IS, recreating the original IS insertion site. This non-replicative mechanism leads to a potential extinction of the IS if transposition is not accomplished ^21^. To prevent this IS loss, IS605 (and IS607) elements use the RNA-guided nuclease TnpB, the likely ancestor to Cas12. TnpB records the sequence of the right end flanking region in its guide RNA and in a similar manner to Cas12, utilizes the guide to programmably cleave specific sequences. Like the PAM of Cas12, TnpB requires an additional Target Adjacent Motif (TAM) to enable cleavage, that is the same as TnpA’s preferred insertion motif. Upon IS excision the resulting scar brings together the TAM and flanking region to produce the ideal substrate for TnpB cleavage. The resulting dsDNA break triggers homologous recombination to copy the IS to the now IS-daughter strand to ensure the IS is never lost from the specific insertion site, akin to homing endonucleases ^22^. Functionally, TnpB records the prior insertion site of the IS then cleaves any such instance it encounters.

To measure the overall impact of TnpB we calculated the total co-occurrence frequency of IS605 and IS200 elements. IS605 had double the co-occurrence frequency compared to IS200 (0.022 for IS200 vs 0.054 for IS605) (**Figure 1G**). This increase in co-occurrence persisted when correcting for multiple potential sources of bias. The PLSDB database is derived from publicly available genomes on NCBI and is skewed towards *E. coli*, where the majority TnpB occurred (**Supplemental Figure 2A**). The trend was maintained when eliminating *E. coli* and highly similar plasmid sequences (**Supplemental Figure 2B**). To isolate potential strain effects, we repeated the analysis on all genomes from *E. faecium* due to the high density of MGEs and found the same trend (**Supplemental Figure 2A/B)**. The effect was also independent from any evolutionary bias of TnpA (**Supplemental Figure 2C)**. Our results indicate that the addition of TnpB fundamentally changes the interaction between the IS and plasmids, biasing the fixation rate to increase overall co-occurrence of the IS in plasmids.

**Figure 2.**
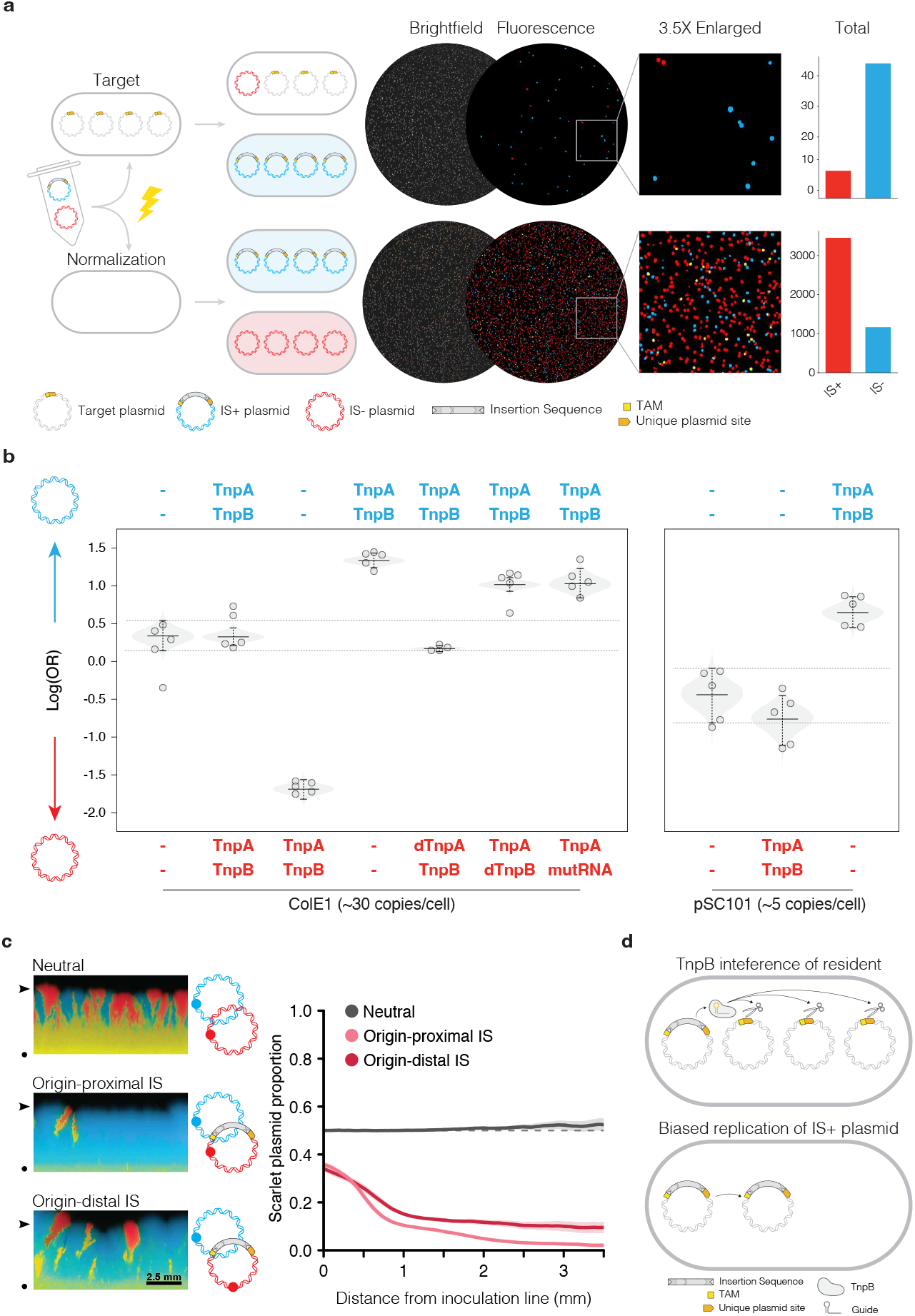
TnpB enables IS+ plasmids to more efficiently displace IS-plasmids. A) Experimental workflow for plasmid displacement assay and example replicates. Fluorescent images are representations of the fluorescence signal that have been enhanced for visualization. The TAM (yellow) and unique plasmid site (orange) together make up the matched target for TnpB. B) Log-odds ratio of colony counts for ColE1 plasmid displacement assays. mWatermelon plasmid variants are labeled on top and mScarlet on bottom. Individual replicates are plotted with the expected posterior in light grey and 95% confidence intervals. Dotted line is 95% confidence interval from the empty-empty plasmid condition. The right box contains selected data for pSC101 origin plasmids. C) Plasmid competition between IS+ and IS-plasmids. IS+ plasmids are highly outcompeted by IS-regardless of placement on the plasmid. The circle indicates the inoculation line and the arrow the final distance measured. D) Proposed gene drive model for the advantage of IS+ plasmids. Insertion sites/TAM is indicated in yellow with the genomic flanking region in green and plasmid site in orange.

### TnpB functions as a gene drive to allow IS+ plasmids to overcome stochastic fixation

To examine our observation that TnpB is beneficial for plasmid occupancy we sought to experimentally measure whether a plasmid containing an active IS605 had an advantage in replacing IS-negative copies of itself. We developed a plasmid displacement assay where a mixture of two fluorescently labeled plasmids are electroporated into a test cell line harboring a non-fluorescent target plasmid (**Figure 2A**) with a validated insertion/excision site from an IS605 ^23,24^. By counting the resulting fluorescent colonies, we can measure the relative advantage of target plasmid replacement by the two labeled plasmids. For example, by adding an IS605 to the blue plasmid and leaving the red empty we can measure the advantage IS605 provides to a plasmid in replacing the resident compared to no IS (**Figure 2A**). In parallel, the plasmid mix was electroporated into plasmid negative cells as a normalization control. From fluorescent images we counted colonies and compared the ratio of IS+ vs IS-colonies in the test and normalization plates to determine the log odds ratio (**Supplemental Figure 3A/B**), reflective of the each plasmids comparative ability to replace the resident (defined as positive if the mWatermelon plasmid had the advantage and negative if mScarlet did). Introduction of a plasmid containing an IS605 with a matched insertion site to the target plasmid containing cells mimics a transposition event into one plasmid within a population, in which one member of an otherwise isogenic population gains a copy of the IS.

**Figure 3.**
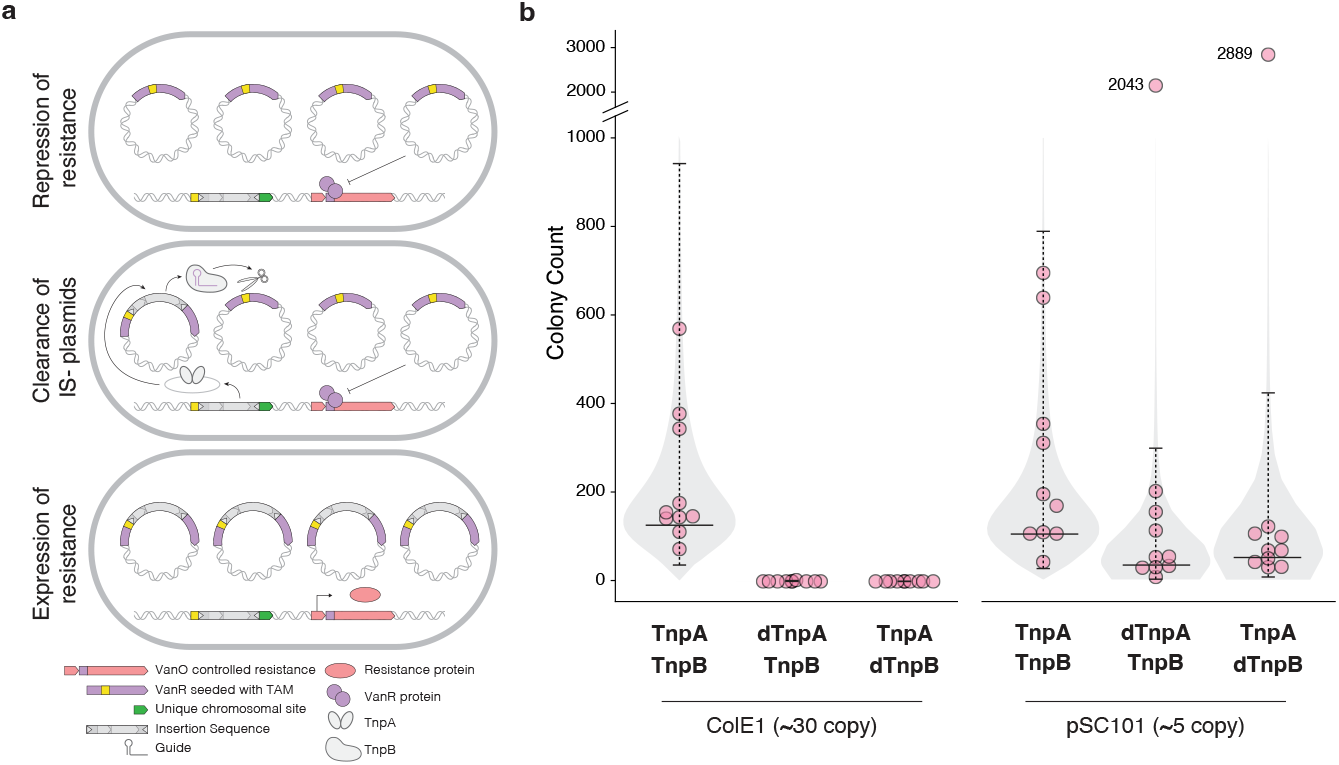
TnpA/B ISs are captured at higher rates than TnpA ISs in a plasmid targeted transposon trap. A) Experimental workflow for transposon trap assay. Vanillin repressor in purple, resistance cassettes in pink, transposon in grey. Yellow indicates potential insertion sites for TnpA. B) colony counting results for insertion sequence variants and target plasmid pairs (indicated on bottom). 95% confidence intervals are indicated with bars around the median from 10 individual replicates as determined from an expectation maximization of the Luria-Delbrück distribution. Outliers are indicated with their colony counts above the dotted line.

IS605+ ColE1 plasmids displayed a significant advantage displacing the target plasmid compared to IS605-plasmids, consistent with our bioinformatic predictions (**Figure 2B**). Matched plasmids, either both IS605+ or IS605-, produced near zero log odds ratio indicating neither plasmid had an advantage and the assay was functioning as expected (**Figure 2B**). Importantly, this advantage was not due to a copy number difference between plasmids as all ColE1 variants had similar plasmid copy numbers from qPCR measurements (**Supplemental Figure 3C**).

The presence of IS605 on its own was detrimental to intracellular plasmid competition. In a within cell plasmid competition experiment plasmids containing an IS605 were strongly outcompeted by those without, regardless of the IS location (**Figure 3C**) ^15^. The disadvantage is likely caused by the added transcription imposed by the IS, inhibiting plasmid replication as recently described ^15^. Thus, the advantage seen in our plasmid displacement experiment was not due to a fitness advantage provided to the plasmid by IS605. Moreover, the results in figure 2B are significant despite the large detriment of the IS to intracellular plasmid competition.

TnpB’s nuclease activity and RNA guide are responsible for the displacement advantage. Catalytic inactivation of TnpA (dTnpA) did not greatly affect the ability of a plasmid to replace the target, indicating TnpA, and subsequent transposition activity, is not responsible for the advantage. TnpB cleaves dsDNA through its RuvC domain which can be inactivated by a point mutation to create a dead TnpB (dTnpB) ^23,25^. Competition of two IS605+ plasmids but with one harboring a dTnpB resulted in a log-odds favoring the unmodified IS605, thus dTnpB removed the advantage provided by IS605 (**Figure 2B**). As the RuvC domain is required for DNA cleavage, the nuclease activity of TnpB is likely driving the observed difference. Moreover, scrambling the guide region of the IS605 (mutRNA) to prevent recognition of the target sequence produced a similar result with the defective guide RNA plasmid losing any replacement advantage.

The advantage provided by TnpB is maintained over different multicopy plasmid types. We repeated the plasmid displacement experiment with the lower copy number pSC101 origin (∼5 copies per cell compared to ColE1’s ∼30) and produced a similar result with IS605 plasmids having a displacement advantage compared to IS605-(**Figure 2B)**. A lower copy number would increase the rate of stochastic fixation for any novel plasmid, thus any advantage given to a low copy number plasmid in our assay would be predicted to have a lessened effect. We did observe a diminished magnitude of TnpB advantage with the pSC101 origin confirming that TnpB’s effect is maintained over different replicons and is sensitive to copy number.

As further support that the IS605 plasmids were actively displacing the target plasmid population we observed diminished fluorescence of IS605-plasmids in test plates. Total fluorescence in the normalization plates was similar regardless of IS605 presence, suggesting the IS does not affect plasmid copy number which could account for the observed advantage (**Supplemental Figure 3D**). In the target plates however, IS605+ plasmids resulted in brighter colonies than IS605-indicating the proportion of IS605+ fluorescent plasmids expands within the cell and thus colonies reflective of an increased fixation rate (**Supplemental Figure 3D**).

The results of our displacement experiment are largely insensitive to image thresholding effects although there exists a slight bias towards mWatermelon (**Figure 2B, Supplemental Fig 3E**). We attribute the mWatermelon bias to a greater fluorescence intensity in our equipment, as exemplified in the double negative condition where the bias is most magnified as brighter colonies are easier to detect. However, this did not affect the results as the IS+ plasmid conditions were insensitive to thresholding effects (**Supplemental Fig 3E**).

We conclude that TnpB’s RNA-guided nuclease activity enables the effective displacement of the resident plasmid population and results in an increased fixation rate. We propose that the programmability of TnpB’s guide enables a specific mechanism of the IS to fix within the plasmid population through a gene drive. When the IS inserts into a plasmid, TnpB will be reprogrammed to the unique plasmid insertion site. Since plasmids are initially genetically identical, this insertion site will exist on all other plasmid copies (**Figure 2D**), providing a guide matched substrate for TnpB cleavage. Cleavage by TnpB will result in destruction of a plasmid and bias the replication of IS+ plasmids, regardless of any fitness disadvantage the IS may impose on the plasmid.

The described displacement experiment mimics a transposition event with a target plasmid designed to specifically match the TnpB guide in the IS. This design allows isolation of the variables to affect displacement advantage but does not incorporate reprogramming of TnpB’s guide nor the transposition activity of TnpA. To test whether this ability is enabled by TnpB’s programmable RNA guide, we sought to evaluate whether this effect holds when an existing plasmid gains a new IS.

### Fixation of novel transpositions into plasmids is enabled by TnpB

To measure the effect of new insertion events in plasmids we devised a transposon trap like those designed to discover novel transposon activity ^26^. We integrated an IS607 with validated high levels of transposition ^27^ and active TnpB into the chromosome along with a resistance cassette under the control of the vanillin operator. We then introduced a target plasmid expressing the vanillin repressor into the cells, resulting in repression of the chromosomal resistance gene and preventing survival on selective media (**Figure 3A**). We designed the repressor coding region to be enriched with potential TnpA insertion sites, i.e. TAMs, setting up a circuit where transposition by TnpA would disrupt the repressor coding region, producing a non-functional repressor (**Figure 3A**). However, so long as a single copy of an intact repressor plasmid exists enough protein will be produced to effectively repress the resistance, preventing survival. Only when a disrupted repressor plasmid fixes within a cell will no repressor be synthesized, enabling survival. By counting colonies on selective plates, we can quantify transposition-and-fixation rates. As mutations in the operator could cheat the circuit, we incorporated fluorescent reporters to mitigate non-IS mediated mutation events (**Supplemental Figure 4 TBS**)

**Figure 4.**
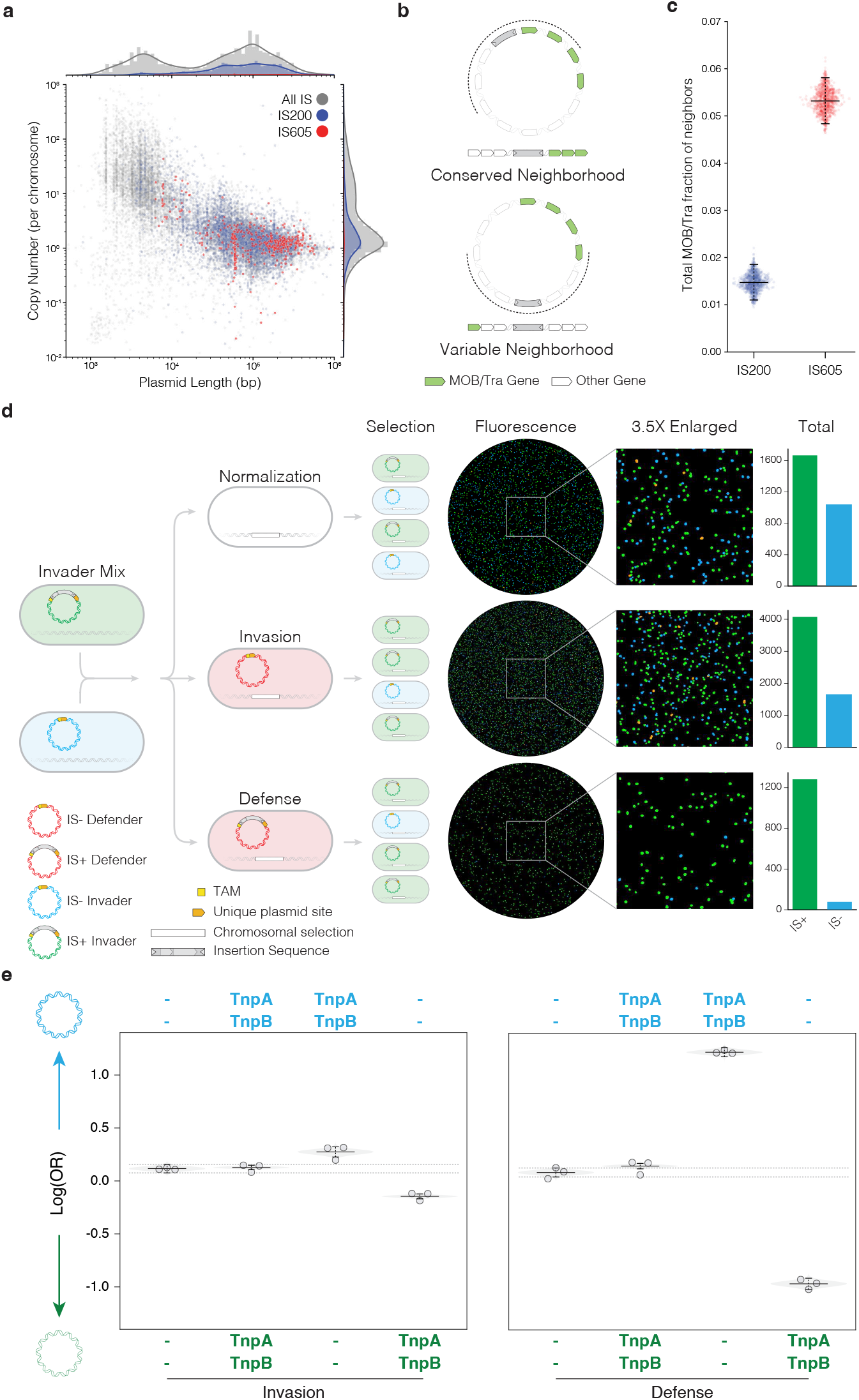
Invasion and defense of conjugative plasmids is benefitted by the presence of TnpB. A) Copy number vs length of plasmids from PLSDB. Short read sequencing was downloaded and aligned to the reference genomes for each available plasmid. The coverage of the chromosome and each plasmid was used to determine the copy number. B) Neighborhood analysis of IS in plasmids and examples of a conserved (top) and variable (bottom) neighborhood around an IS. For each plasmid in the PLSDB database the transfer and mobilization genes (Tra/MOB genes) are identified by MOB-suite (green). For each gene neighborhood of an IS (indicated by the black arch) the fraction of MOB genes is determined. The closer the IS is to the transfer and mobilization operons, the greater the fraction of MOB genes in the neighborhood and the conservation of IS location. C) Total fraction of MOB neighbors for IS200 and IS605 in the PLSDB dataset. Error bars are 95% confidence intervals determined from N=1000 bootstrapping of the data. D) Experimental set up for conjugation replacement assay. E) Log-odds quantification of invasion (left) and defense (right) conditions from the conjugation invasion assay with the *E. coli* W1485 F plasmid.

An active IS in the chromosome produced significant survival after incubation and selection with a ColE1 target plasmid in the transposon trap (**Figure 3B, left**). The result of 10 individual replicates followed an expected long tailed Luria-Delbrück distribution, common to similar mutation accumulation experiments ^28^. Catalytic inactivation of TnpA resulted in extremely minimal survival, which was expected as the inability to transpose would prevent any disruption of the repressor and completion of the circuit (**Figure 3B, left**). Together these two results indicate the circuit works effectively and any decrease in survival would be indicative of an impaired ability to disrupt the plasmid population.

Inactivation of TnpB resulted in no detectable survival in the ColE1 condition, demonstrating that the programmable nuclease ability of TnpB was critical to fixing the plasmid population (**Figure 3B**). We repeated the experiment with the lower copy pSC101 origin in the target plasmid and found a similar level of survival with an active IS compared to ColE1. However, for the dTnpA and dTnpB conditions we observed detectable levels of survival with some replicates within the range of the fully active condition (**Figure 3B, right**). As with the superinfection experiment, we expected the low copy number to decrease the relative importance of the effect of TnpB. The low copy number of pSC101 allows for a greater stochastic advantage of any novel plasmid, whether that be from an insertion event or a mutation. Some survival in the dTnpA and dTnpB conditions was likely due to background mutations that we could not eliminate. There was a slight increase in survival for the dTnpB condition which we did expect to see as the transposase is still active and able to disrupt the repressor. The pSC101 result demonstrates that the effect of TnpB is magnified with the increased copy number of the target plasmid. Taken together, the transposon trap reveals that TnpB improves fixation of newly-introduced IS in multicopy plasmids, to the point of being effectively required for higher copy plasmids.

### TnpB provides an advantage to intercellular competition of conjugative plasmids

While we find that the strength of the TnpB gene is increased with a higher copy number (**Figure 2B**), ISs are much more common in larger plasmids, likely due to the increased cargo capacity (**Supplemental Figure 5A**) ^4,5^. We determined the copy number relative to the chromosome of all plasmids with short read data in the PLSDB dataset and found that indeed IS200 and IS605 elements tended to be found in large, low copy plasmids (**Supplemental Figure 5B, Figure 4A**). We hypothesized that the location of IS605 elements on plasmids may not be random as TnpB’s targeting mechanism depends on the insertion site; the specific location could enable targeting or other plasmids. As low copy plasmids are often conjugative, coding for machinery to enable HGT, we moreover posited that TnpB could provide an intercellular advantage.

TnpB-containing ISs clustered in conserved plasmid regions. We identified transfer and mobilization genes (MOB) with MOB-suite to conduct a neighborhood analysis of IS200 vs IS605 elements in plasmids as a proxy for an IS residing in a conserved plasmid region (**Supplemental Figure 5C**) as MOB genes are highly conserved between plasmids ^29^ (**Figure 4B**). IS605 elements had a significantly increased total fraction of MOB gene neighbors, strongly suggesting that IS605 clustered in more conserved plasmid regions, enabling TnpB to target more conserved genetic sequences (**Figure 4C**). This enrichment held across a range of neighborhood windows (**Supplemental Figure 5D**) and was not due to any evolutionary bias (**Supplemental Figure 5E/F)**. While we did not identify the specific TnpB guides to determine targeting potential, our results strongly suggest that it is possible for TnpB to target a range of plasmids through residence in a conserved region.

We created a system to measure competitive invasion advantage of conjugative plasmids. Similar to the superinfection experiment we integrated fluorescent labels (sfGFP and mTurquoise2) onto the F plasmid of *E. coli* with either an IS605 or TnpB target site to create invader strains. We created a recipient strain with a potential TnpB target site within the F plasmid along with a fluorescent marker and a chromosomally integrated resistance cassette to select for transconjugants (**Figure 4D**). A mixture of two invader strains was incubated with the target for 24 hours before selecting for transconjugants and taking fluorescent images (**Supplemental Figure 6A)**. At the same time the invader mix was incubated with empty for normalization enabling quantification of the advantage to invasion granted by the IS through a similar log odds ratio as the displacement experiment (**Figure 4D, middle**). As a plasmid containing an IS could also be the target of invasion, we also swapped the resident target plasmid for one containing the IS. Thus, the assay enables us to measure the defensive advantage an IS grants a conjugative plasmid in addition to the invasion benefit (**Figure 4D, bottom**).

IS605 provided an advantage to both plasmid invasion and defense. Competition of matched invader strains (both with or without IS605) produced near zero log odds in both experimental conditions as expected (**Figure 4E**). IS605 provided a modest, but significant advantage in plasmid invasion (**Figure 4E, left**), indicating TnpB was clearing the resident target plasmid to enable invasion. When selecting for superinfected cells, those where the invader and target plasmid were both present (**Supplementary figure 6B**), we only observed colonies where the invading TnpB was catalytically inactive, further supporting that TnpB was cleaving the target plasmid (**Supplementary figure 6C**).

In the defense condition IS605 provided a much larger advantage (**Figure 4E, right**). We reasoned the difference in magnitude between invasion and defense to be a result of differing conjugation and transcription kinetics in relation to genetic drift. For the advantage to be realized in the invasion condition the invader plasmid must enter as ssDNA, re-circularize then transcribe TnpB, a relatively slow process compared to the genetic drift of low copy plasmids in actively dividing cells. However, in the defense scenario TnpB is constantly transcribed by the resident cells, able to immediately cleave the invading plasmid, analogous to the function of CRISPR. Given recent work on the leading region of conjugative plasmids and TnpB’s ability to target ssDNA, we propose that the orientation and location of the IS could further modulate the strength of the defensive advantage ^25,30^.

## Discussion

In this work we uncovered a positive association between IS chromosomal copy number and plasmid co-occurrence across the majority of IS families (**Figure 1D-E, Supplementary Figure 1C**). This association points towards two complementary factors favoring the emergence of IS-bearing plasmids. First, high IS chromosomal copy number might provide greater transposition opportunities resulting in increased transposition rates to plasmids. Second, high IS chromosomal copy number may be indicative of a more permissive cellular environment to transposition, such as decreased regulation. However, a few IS families deviated from this general trend, suggesting mechanisms beyond a numerical advantage to increased plasmid carriage. Plasmid-specific mechanisms of IS invasion could fulfill a powerful role in the spread of their ISs across bacterial populations by facilitating the ability of ISs to rapidly increase their prevalence, and hitchhike on mobile plasmids. In searching for such a mechanism, we focused on the closely related IS200 and IS605 families that displayed distinct patterns of association with plasmids. The co-occurrence of IS200 in plasmids was strongly dependent on high chromosomal copy number. Yet IS605 elements, which are distinguished from IS200 by the presence of single additional protein TnpB, co-occurred in plasmids much more frequently, particularly at low chromosomal copy number, breaking the general trend. Therefore, we hypothesized that TnpB could be enabling a specific mechanism favoring the acquisition or spread of ISs in plasmids.

We found that TnpB facilitates the invasion of plasmid populations via a unique mechanism. The RNA guided nuclease activity of TnpB was originally characterized as promoting chromosomal IS retention by cleaving the excision scar left from transposition ^22^. Our work revealed a complementary but distinct role for the programmable ability of TnpB’s nuclease activity. In the context of plasmids, a nascent transposition event results in the reprogramming of TnpB’s guide to target IS-plasmids, effectively biasing replication and inheritance towards plasmids that are IS+ (**Figure 2B, Figure 3B**). This biased inheritance is the defining feature of a gene drive. Although usually considered a phenomenon tied to eukaryotic sexual reproduction, gene drives in principle only require a high ploidy cellular environment. Our findings showcase that even the simplest instances of high ploidy, such as plasmids, can support naturally occurring gene drives.

Plasmid gene drives display complexities absent from their eukaryotic counterparts. While eukaryotic gene drives usually occur in diploids with one chromosome acting as the driver and its homolog the target, plasmid gene drives occur in truly polyploid environments with multiple targets. The existence of plasmids in varying copy numbers allows us to investigate the role of varying target numbers on the gene drive. We found that the higher the plasmid copy number (i.e. ploidy), the greater the effect of the gene drive (**Figure 2B**), in line with theoretical predictions on multi-scale evolution ^31^. This observation can be understood by the lesser impact of stochastic fixation with higher copy number and the consequently increased effect of the gene drive. However, a refinement of our method is required to confirm the exact modulation effect by copy number, as our results are potentially confounded by protein dilution effects particularly with the transposon trap. Nevertheless, plasmid copy number is a key factor in the success of the IS invasion. Overall, the emergence of an IS+ plasmid is not simply the result of a transposition event, but part of a complex interaction network that is jointly modulated by plasmid ploidy and IS chromosomal copy number.

The plasmid gene drive mechanism provides a plausible explanation for the broad distribution of TnpB itself. TnpB is a common protein occurring in roughly 30% of all bacterial genomes, despite their respective ISs having lower predicted mobility than other IS families^32,33^. The described gene drive could have promoted the spread of TnpB in two distinct scales. First, by favoring the initial emergence of IS+ plasmids harboring TnpB (**Figure 3**). Second, by providing IS+ conjugative plasmids with invasion and defense advantages against similar IS-plasmids competing for the same hosts (**Figure 4**). The enhanced spread of TnpB across genomes has likely enabled the functional diversification of these proteins as transcriptional modulators^34,35^ and most notably as the Cas12 effector in CRISPR^23,32^. In particular, the defense advantage given to conjugative plasmids is analogous to the classically described function of CRISPR, providing protection against other MGEs ^32^.

Previous work detailing the evolutionary relationship between Cas effectors and transposon nucleases has found independent instances of guide RNA accumulation, sometimes even resembling CRISPR arrays ^32,33^. These events were proposed as evolutionary bridges between transposon-bound nucleases and the modern defense CRISPR-Cas systems. Although these observations demonstrate how the structure of CRISPR-Cas systems might have originated, they do not explain how such systems gained their defensive functions. Our conjugation results suggest that at least some defensive functions emerged naturally from the interaction of ISs and plasmids even prior to the development of arrays. In effect, IS integration into plasmids and other MGEs might have provided their nucleases with an ancestral evolutionary path between transposon retention and defense activities.

While TnpB’s gene drive mechanism has likely benefitted the spread of TnpB itself, ISs broadly enable the spread of genes including those outside the bounds of the IS. ISs frequently serve as recombination hot-spots to allow mobilization of normally immobile elements, enable adaptation of bacteria to environments, and allow plasmid reorganization of genes through composite transposons ^36–38^. Antimicrobial resistance (AMR) is particularly subject to spread by plasmids and is heavily modulated by ISs that serve to distribute AMR genes across plasmids ^8,9,39^. Absent from the understanding of the role ISs play in AMR acquisition by plasmids, however, is how an IS comes to reside on the plasmid in the first place. For AMR to spread through plasmids via an IS, a plasmid must first acquire the IS itself. The spread of AMR on plasmids is thus a consequence of IS presence, but not the reason for their existence. TnpB’s promotion of IS incorporation into plasmids is a specific mechanism to achieve IS presence on a plasmid, resulting in potentially powerful subsequent diversification from incorporation of new genetic material besides the IS itself.

Plasmids steer bacterial evolution both by direct fitness effects and indirect evolvability effects. Typically, the spread of genes in bacteria is associated with direct, first-order host-advantageous traits such as AMR. In contrast, the acquisition of ISs by plasmids is a case where gene dissemination regularly occurs in the absence of any direct host fitness benefits. Despite this lack of a first-order host advantage, IS acquisition provides critical second-order benefits to hosts by enabling gene exchanges across the bacterial pangenome and boosting the evolutionary flexibility of the host cells. However, it is generally unclear to which extent second-order effects actively drive the fixation of novel variants in the absence of a first-order benefit, as is the case for IS-bearing plasmids ^12,14^. Here we have shown that within-cell dynamics can drive the fixation of some classes of IS-bearing plasmids, bypassing the need for first-order fitness benefits and empowering the evolvability of host cells. This unexpected connection between multi-scale selection and the evolvability of bacteria highlights the importance of the nonlinearities introduced by MGE interactions. Crucially, IS-bearing plasmids spread beyond the ability of either MGE individually. Far from being passive passengers on other MGEs, ISs can encode active mechanisms to bias their carriage and spread. We argue that the nested existence and subsequent interactions between MGEs are not an incidental result of selfish spread, but a driver of it.

## Acknowledgements

We thank Ákos Nyerges and the laboratory of George Church for supplying the MDS42 cells, Arya Kaul, Siân Owen, Suchita Nety and Soumya Kannan for helpful discussions. This work was supported by the NIGMS of the National Institutes of Health (R35GM133700 and R35GM156320), the Pew Charitable Trusts and NSF grant MCB2426105. K.S.M. was supported by an NDSEG Fellowship.

