## Supplemental Figures for "RNA-guided nucleases enable a gene drive of insertion sequences in plasmids"

### Materials and Methods

*Curation of PLSDB database*

The PLSDB meta archive and triangle file was downloaded from https://ccb-microbe.cs.uni-saarland.de/plsdb2025/download then unique assembly NCBI accession numbers collected. Assembly genbank and FASTA files were downloaded from NCBI and replicon accession numbers and identities added to our database metadata. Assemblies that did not have genbank annotations were annotated with Prokka. Highly similar plasmids were collected into families from PLSDB's hash table to identify redundant sequences.

*Insertion sequence identification*

ISEScan was run on each assembly and the output collected into a single metadata file with a custom python script. IS presence on plasmids vs chromosomes was determined from the ISEScan results and PLSDB metadata. The IS family and plasmid/chromosomal copy number was determined from the ISEScan output [19].

To find specific IS200, IS605 and IS607 subgroups a HMMER search was performed separately on the PLSDB genbank files using models NF033573.1 and NF033518.0 for TnpA and models TIGR01766.2, NF040570.1, NF038281.1, NF038280.1, PF07282.15, NF040563.1 for TnpB/IscB. The final HMMER output was collected into a single file and pairs of TnpA and TnpB identified as those being within 1000 kb of each other with a custom script. An E-value threshold of 10E-25 was used to filter the final hits and presence on plasmids and chromosomes identified. A pairwise blast or MMSeqs2 were utilized to identify clusters (sequence identity >90%). Clusters were then used to determine the chromosomal copy number.

*Co-occurrence frequency*

To determine co-occurrence frequency genomes were filtered for IS chromosomal copy number greater than two to ensure the IS was mobile. Each IS cluster in each genome was scored as 0 in the cluster only occurring in the chromosome and 1 if in the chromosome and plasmid. We defined co-occurrence frequency as the total hits (i.e. each IS cluster genome pairing that was scored at 1) divided by the total genomes.

A sigmoid was fit to the raw scored data via a maximum likelihood, equation 1, where p(x) is the predicted co-occurrence frequency at a given chromosomal copy number. $\beta_0$ and $\beta_1$ are constants, the latter being defined at the rate parameter which reflects whether the association is positive or negative. Binned co-occurrence frequencies were calculated by determining the co-occurrence frequency for a window range of chromosomal copy numbers and a beta distribution prior determined with a custom python notebook. The $\beta_1$ for each IS family was extracted from the corresponding sigmoid fit (**equation 1**) and 95% confidence intervals determined by a chi-squared approximation from a profile likelihood of the rate parameter.

$$\text{Equation 1: } p(x) = \frac{1}{1 + e^{-(\beta_0 + \beta_1 x)}}$$

*Fluorescent imaging*

To ensure high quality images synthetic black iron oxide was added to all LB agar plates to produce a black background. Plates were imaged with a custom fluorescent plate imager. Plates are illuminated by colored LEDs with excitation filters (EX), the emitted light passes through emission filters (EM), and images are taken on a Canon EOS camera. The red channel pairs 567 nm LED with 562 nm EX and 641/75 nm EM filters, the green channel pairs 490–515 nm LED with 494 nm EX and 540/50 nm EM filters, and the blue channel pairs 448 nm LED with 438 nm EX and 483/31 nm EM filters. Exposure times varied between 0.5s and 8s depending on the plate and intensity of fluorescent signal.

Channel intensity images were taken of a white background and no filter for exposure times of 1/20-1/30s to normalize spatial differences in fluorescence. Camera settings were kept at an aperture of 10 and ISO of 200. Final images were intensity normalized using these background images.

*Electroporations superinfection*

TOP10 cells were transformed with a target plasmid containing a dead fluorophore with kanamycin resistance and a TnpB target site for the IsDra2 insertion sequence [23]. Electrocompetent cells were generated by growing to OD = 0.8 at 30 °C, centrifuged at 2500xg, washed with ice cold water before being resuspended in 25% glycerol at 100X concentration. Plasmids 100 ng of each of two plasmids of interest were mixed and diluted to 5uL which were then electroporated into 50uL of target plasmid cells and empty standard TOP10 cells. TOP10 were incubated at 37 °C for 1 hour prior to plating while the target plasmid cells were plated immediately. The same kanamycin resistance as the target plasmid was used in the plasmids of interest, requiring high density plating to obtain enough colonies. This was done to capture fluorescent dynamics, in addition to preventing selection for electroporation events as opposed to fixation events.

Agar plates were prepared with 25mL 2% LB agar, 1% MARS synthetic iron oxide plates and kanamycin. A dilution series was prepared of the electroporated cells and 0.5mL of each dilution was mixed with 5mL of 50C 0.5% LB Agar then plated. After allowing the top agar to solidify a second layer of top agar was added. Plates were incubated at 37 °C overnight. The following day the top agar layer was dried at 37 °C until collapsed to allow oxygen mediated maturation of the fluorophores. Plates were incubated at 4 °C overnight then imaged the next day.

Final images were taken and analyzed with a custom python script. Images were first intensity normalized and a mask to eliminate the plastic plate boundary that produced significant autofluorescence in the green channel. A minimum threshold was estimated from the pixel distribution and a watershed segmentation algorithm applied to identify and count colonies. Additional thresholds based on size and intensity were applied then the overall intensity value adjusted to match the green and red channels. Re-coloring and dilation was applied separately for visualization.

*Plasmid displacement assay*

Intracellular plasmid competition experiments were performed as previously described [15]. In brief, a plasmid dimer consisting of a mScarlet labeled plasmid with the IsDra2 transposon with catalytically inactivated TnpA and TnpB, and a mWatermelon labeled plasmid was introduced to cells containing the patagonian FLP and a chloramphenicol resistance cassette Plates with 40mL of 2% LB agar with 1% MARS synthetic iron oxide and 0.3% arabinose were prepared. Inoculation of cells was performed by wetting a razor blade with a saturated cell culture. Inoculated plates were grown for 12 hours at 37°C (in which FLP is inactive) and then transferred to a moist incubator at 30°C for up to 2 weeks. Imaging was performed as described above. Analysis was performed with a custom R script.

*Conjugation invasion*

For the invasion cell lines a cassette containing either the IsDra2 IS or the target site, mTurquoise2 or GFP and chloramphenicol resistance was integrated with lambda red recombinase onto the F plasmid of the W1485 strain *E. Coli*. For the recipient cell lines a similar cassette containing either the IsDra2 IS or the target site, mScarlet and kanamycin resistance was integrated with lambda red recombinase onto the F plasmid of the W1485 strain *E. Coli*, then the engineered F plasmid conjugated into an MG1655 strain with a chromosomal ampicillin resistance gene.

A single colony from the invader and recipient strains was grown overnight at 37 °C in a shaking incubator under selection, then back diluted 1:100 the next day and grown without selection for 2 hours or until an OD600 of 0.5. The two invader strains and recipient strain were mixed in a ratio of 1:1:8 to prevent cross conjugation of the invaders, the diluted 1:10,000 and incubated overnight at 37 °C in a shaking incubator with no antibiotic. The following day the conjugation cultures were diluted, added to 0.5% LB agar top agar and plated on selective (ampicillin and chloramphenicol), non-selective (chloramphenicol) or superinfection plates (ampicillin, chloramphenicol and kanamycin) 2% LB agar plates with 1% synthetic iron oxide. A second layer of 0.5% LB agar top agar was then added then the plates incubated at 37 °C overnight. The following day the plates were dried at 37 °C to collapse the top agar then incubator overnight at 4 °C before being imaged.

Colonies were counted with a custom python script. The fluorescent channels were intensity normalized before a binary mask applied to identify colonies with any fluorescence then a watershed algorithm applied to identify colonies bounds. The fluorescence of each channel within a colony was quantified and threshholded to determine the ratio of GFP to mTurquoise2 colonies. Re-coloring and dilation was applied separately for visualization.

*Transposon trap*

A chloramphenicol resistance cassette and the red reporter mScarlet-I, both with expression controlled by the vanillin operator, were introduced in the LacI locus of the genome of transposon and insertion sequence devoid MDS42 cells [40] with lambda red recombinase. The recently described *Clostridium botulinum* IS607 highly active TnpA and TnpB was introduced into the chromosome at the araBAD locus [27].

We created an optimized coding sequence for the vanillin repressor (VanR) that was saturated with TAMs for IS607 (VanTAM). Plasmids (PBR origin ~20 copies per cell and PSC origin ~5 copies per cell) carrying VanTAM, the green reporter protein mWatermelon, and a Kan resistance cassette were transformed into these cells. A dilution series was performed and inoculated into 1mL in 96 well-plates. Cultures were grown to saturation for 18 hours. Dilutions in which each well was inoculated with on average at most a single cell were selected by checking that most wells were empty after the growth period. The content of ten wells that had cultures grow to saturation was mixed into 3mL of 0.5% agar and overlaid on dishes containing Kan and Chlor. In this way, only cells that retained the plasmid but disrupted the repressor could form colonies. Moreover, the green marker allowed us to assess that the plasmid was retained, and the red marker functioned as an independent marker for the disruption of the repressor gene. Plate imaging was performed as in the other assays and colonies counted with a custom python script based on a binary mask and watershed algorithm. Only colonies that were both green and red were counted.

*qPCR for plasmid copy number estimation*

qPCR was used to determine the relative plasmid copy number per chromosome in each plasmid-containing strain. As unbiased quantification of both chromosomal and plasmid DNA is required for an active copy number estimate, several steps were conducted to mitigate potential sources of bias in both the experimental and analytical techniques. For each strain, gDNA extraction was performed. First, lysate was prepared using 1 mL of bacterial culture in active exponential growth (OD 0.5, N=3) for each strain using Invitrogen PureLink® Genomic Digestion Buffer and Thermolabile Proteinase K (New England Biolabs), followed by an incubation at 55°C for 1 hour and 25 minutes of heat inactivation at 65°C.

In order to prevent size-biased loss of plasmid or gDNA from any column purification steps, gDNA was precipitated with isopropanol. Before beginning precipitation, sodium acetate was added to a final concentration of 0.3 M and well-mixed into the lysate to prevent the SDS in the digestion buffer from interfering with precipitation. After isopropanol was added, lysate was centrifuged for 1 hour at 4°C before the pellet was washed twice with 70% EtOH. Extracted gDNA was then resuspended in molecular grade water before proceeding.

To prevent plasmid supercoiling from interfering with plasmid PCR efficiency relative to the chromosome and confounding copy number estimates, genomic DNA (gDNA) extractions were first digested with the restriction enzyme SalI-HF (New England Biolabs). This enzyme was selected because it cuts the plasmid and chromosome at sites that do not overlap with the qPCR primer targets. Digestions were performed for 1 hour at 37°C, followed by heat inactivation at 65°C for 25 minutes. The digested DNA was used directly as the qPCR template without purification, as it was substantially diluted in the final reaction. Quantitative PCR was performed with Luna® Universal qPCR Master Mix (New England Biolabs), targeting 100 bp regions on both the chromosome and plasmid for each strain. Two template dilutions per replicate were used to allow for downstream calculation of PCR efficiencies.

Differing PCR efficiencies in the target in the plasmid relative to the chromosome greatly confounds calculation of plasmid copy number from qPCR data. To help account for this, PCR efficiencies ($E$) were calculated for the plasmid and the chromosome reaction by comparing two dilutions of gDNA template using **equation 2**:

$$\text{Equation 2: } E_{plasmid} = 10^{1/\Delta q_{plasmid}} - 1 \text{ and } E_{chromosome} = 10^{1/\Delta q_{chromosome}} - 1$$

where $\Delta q$ is the difference in cycle number of the 10-fold dilution. Plasmid copy number relative to chromosome was then calculated using the median PCR efficiency for the chromosomal or plasmid reaction using **equation 3**:

Equation 3: $\frac{(1+E_{chromosome})^{q_{chromosome}}}{(1+E_{plasmid})^{q_{plasmid}}}$ = Plasmid copy number relative to chromosome

Where $q_{chromosome}$ and $q_{plasmid}$ correspond to the cycle number of plasmid or chromosomal reactions happening at the same dilution. Absolute plasmid copy number estimates were then calculated by multiplying this relative value by 3, in order to estimate the approximate number of plasmids in a bacterial cell under active exponential growth.

*Statistical analysis*

For the electroporation and conjugation invasion experiments the log-odds ratio of each condition was calculated. The expected distribution was determined by a Monte Carlo simulation of the aggregated colony counts with a custom python script from which 95% confidence intervals were determined. P-values were determined from a Monte Carlo simulation.

For the transposon trap a custom python script was used to generate the expected distribution of the data from an expectation maximization of the Luria-Delbruck distribution[28,41], from which 95% confidence intervals were extracted.

For the co-occurrence frequency curves the co-occurrence of each binned chromosomal copy number was calculated and the prior beta-distribution determined. A logistic regressor was fit via a log-likelihood expectation maximization and the 95% confidence intervals of the rate parameter determined from a profile likelihood custom python script.

For total co-occurrence frequency comparisons the co-occurrence frequency was calculated and the 95% confidence intervals derived from bootstrapping with N=1000. The same analysis was performed for neighborhood clustering.

*Code availability*

Jupyter notebooks to analyze the data and generate figures can be found in the github repository https://github.com/keplermears-hms/TnpB-gene-drive/tree/main along with bash scripts and snakemake pipelines.

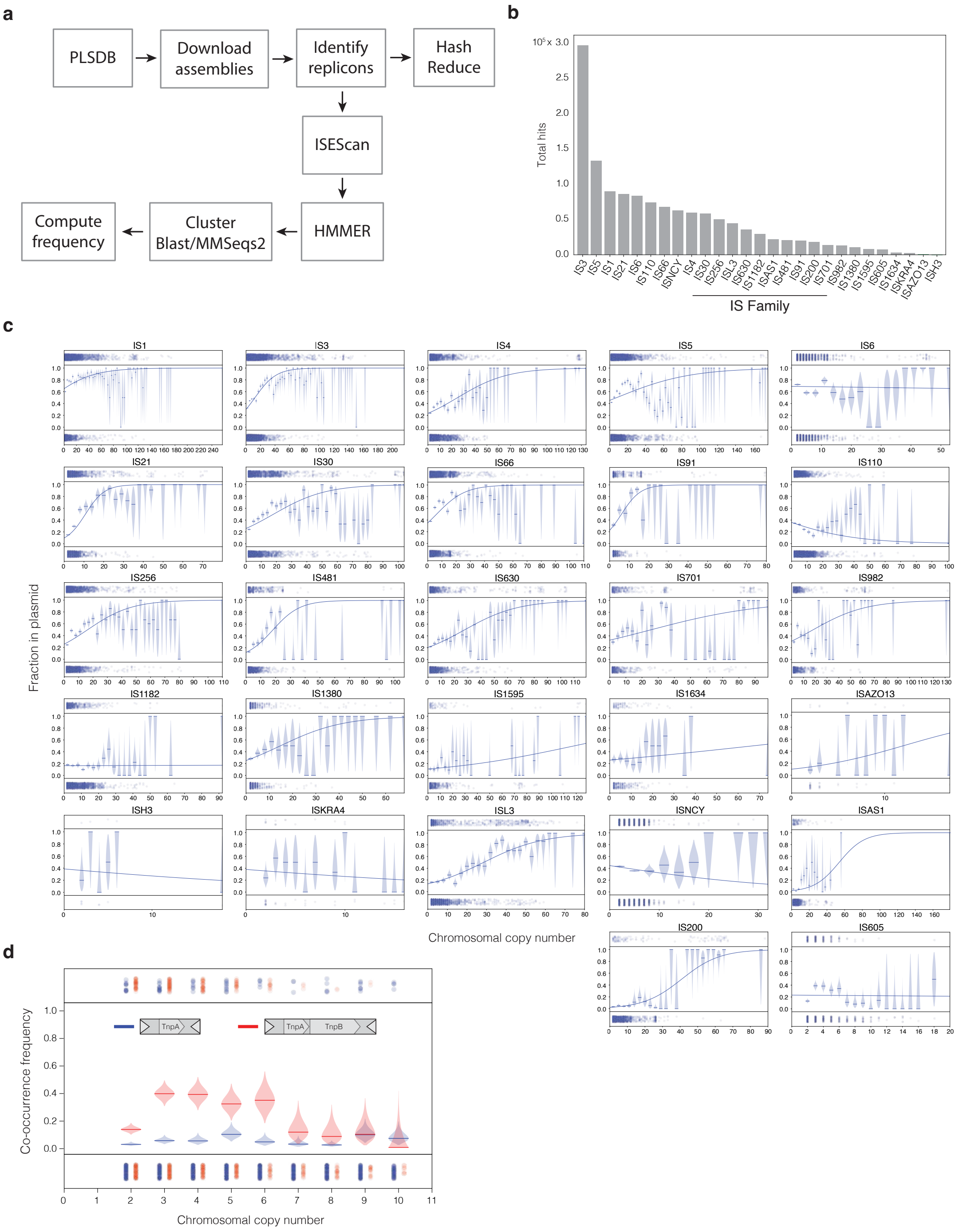


**Supplemental figure 1: Identification of insertion sequence chromosomal copy number and co-occurrence in the PLSDB database**

A) Pipeline for the curation of the PLSDB replicons and identification of insertion sequences. B) Total hits for each IS family from running ISEScan on the PLSDB database. C) Co-occurrence frequency curves and sigmoid fits for all IS analyzed in this study and used to generate rate parameters in figure 1D. D) Overlaid co-occurrence frequencies of IS200 (blue) and IS605 (red) for low chromosomal copy number. At low chromosomal copy number IS605 have significantly higher co-occurrence frequency, in addition to having increased total co-occurrence frequency.

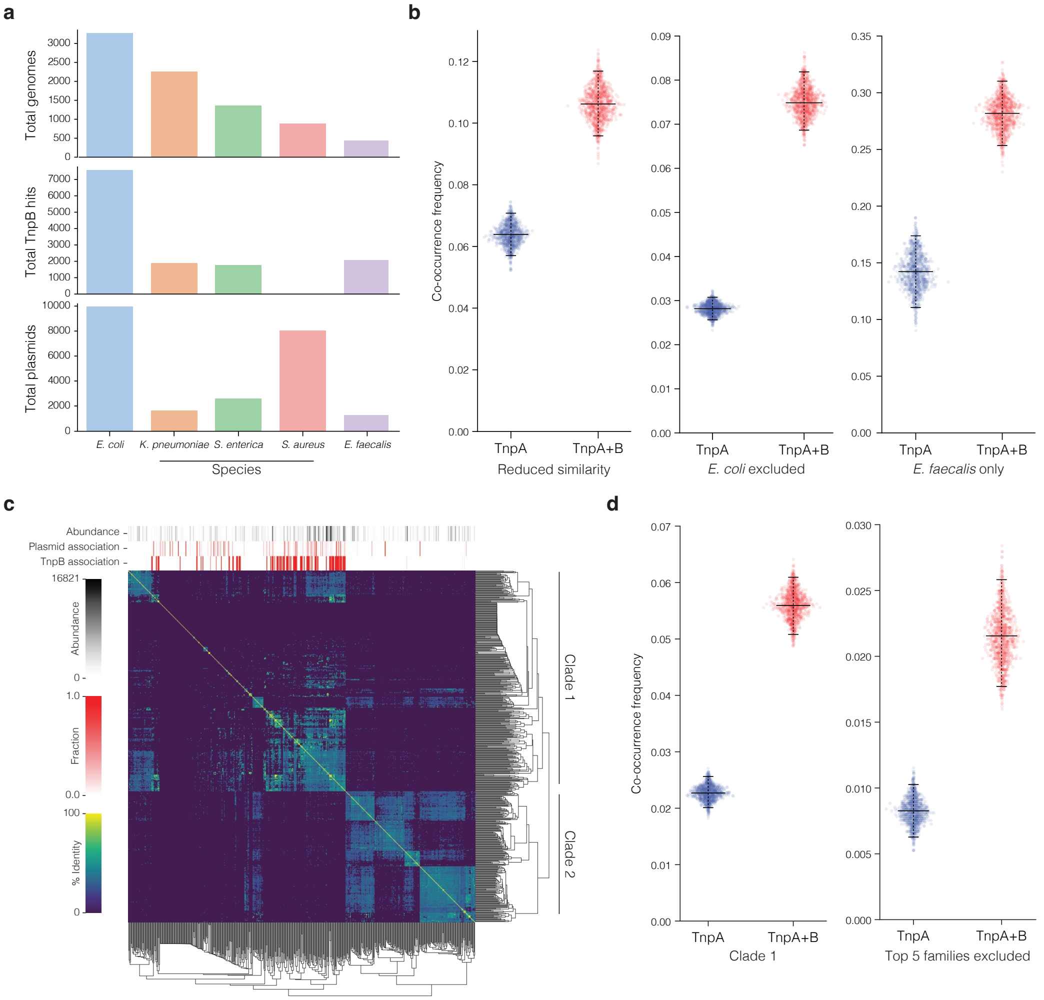


**Supplemental 2: Controlling for bias in IS co-occurrence frequency analysis**
A) Total number of genomes (top), TnpB hits (middle) and plasmids (bottom) for the top five most represented species in the PLSDB database. B) Total co-occurrence frequency for IS200 (blue) and IS605/607 (red) when reducing similarity of plasmids in the PLSDB database (hash reduction using a threshold of 0.02), omitting *Escherichia coli* genomes, and for *Enterococcus Faecium* only. C) Similarity clustering of TnpA from IS200 and IS605 elements. The enrichment of IS605 in plasmids could be due to an evolutionary bias where the divergence of TnpA the two subfamilies is entirely responsible for the effect. TnpAs were extracted and clustered with MMSeqs2 at 50% AA similarity to create families. Representatives from each family were pairwise aligned to create the similarity matrix and the relative abundance, plasmid association and TnpB association determined (top bars) for each family. TnpA families were clustered into two clades, one with greater TnpB association (predominantly IS605/IS607) and one without (IS200). D) Co-occurrence frequencies for IS200 and IS605 from just clade one in (C) and with the top 5 most representative TnpA families excluded (right).

95% confidence intervals for (B) and (C) were determined from bootstrapping (N=1000) with each replicate in light blue/red.

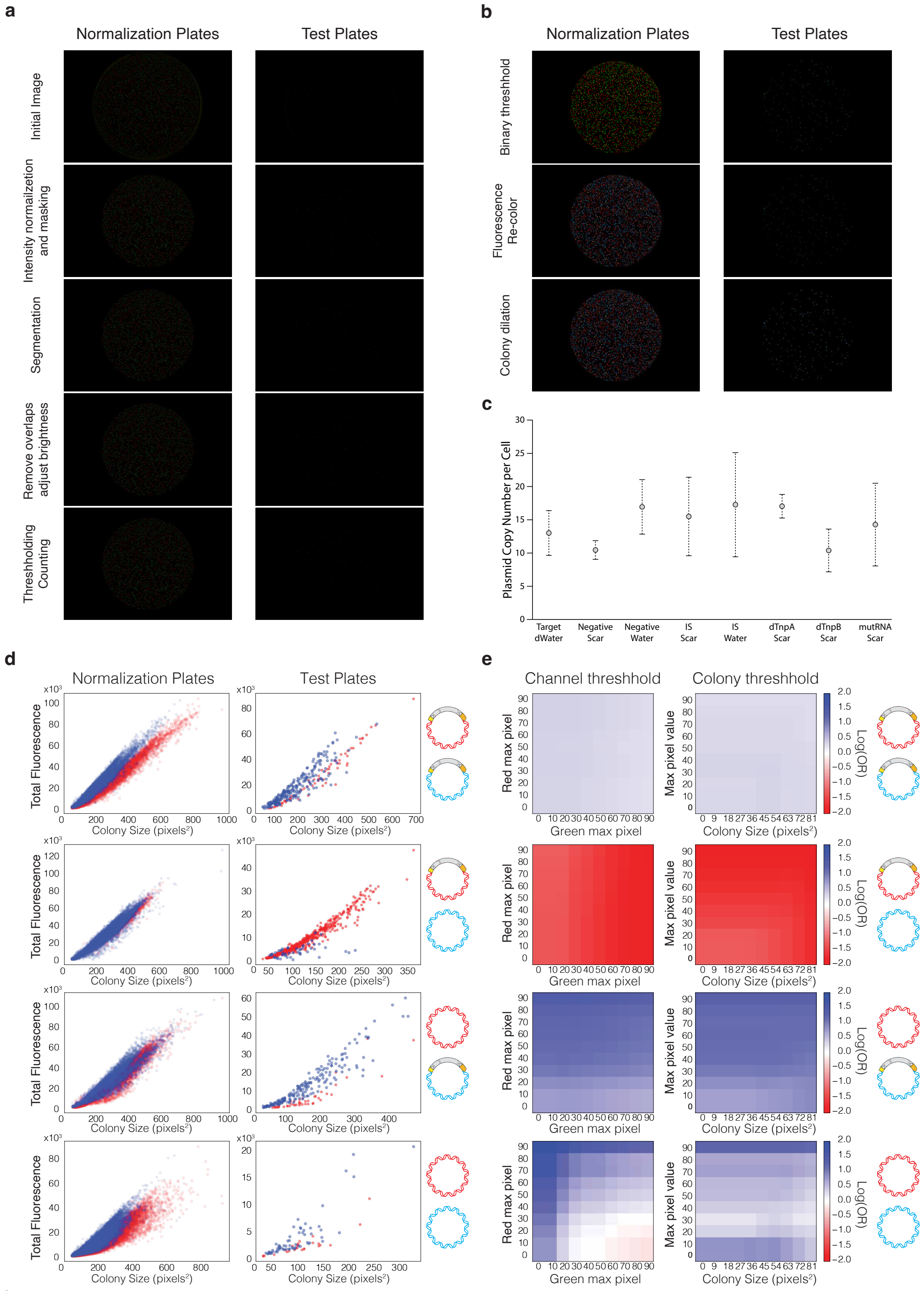


**Supplemental 3: Image analysis pipeline for plasmid displacement assays**

A) Image workflow for colony counting. Initial images were normalized by the relative channel intensity, then masked to only include the interior of the plates. Colonies were identified with a watershed segmentation algorithm then scaled before thresholding applied to count for the analysis. Overlapping colonies are only removed for test plates but counted for normalization plates. B) Image workflow for visualization. Binary threshold was applied to maximize the channel intensities then recolored. Test plate colonies were dilated to appear larger. C) Plasmid copy number per cell for all ColE1 variants in Figure 2B as determined from qPCR. Error bars are standard deviations of 3 replicates and 2 dilutions. D) Colony total fluorescence vs. size of aggregate data for five replicates prior to calculation of the log odds ratio. Plasmid conditions shown on the right in order top to bottom: both with IS, red with IS blue empty, blue with IS red empty, both empty. E) Threshold sensitivity analysis for the conditions in C for individual channel brightness thresholding and overall brightness and size thresholding. Tiling threshold difference in intensity between the fluorescence channels, and in intensity and colony size resulted in minimal deviation of the final log-odds ratio. There was slight sensitivity when competing two empty plasmids, where we saw a bias towards mWatermelon which was mirrored in our results. We attributed this bias to the brighter fluorescence intensity of mWatermelon. Plasmid conditions shown on the right in order top to bottom: both with IS, red with IS blue empty, blue with IS red empty, both empty.

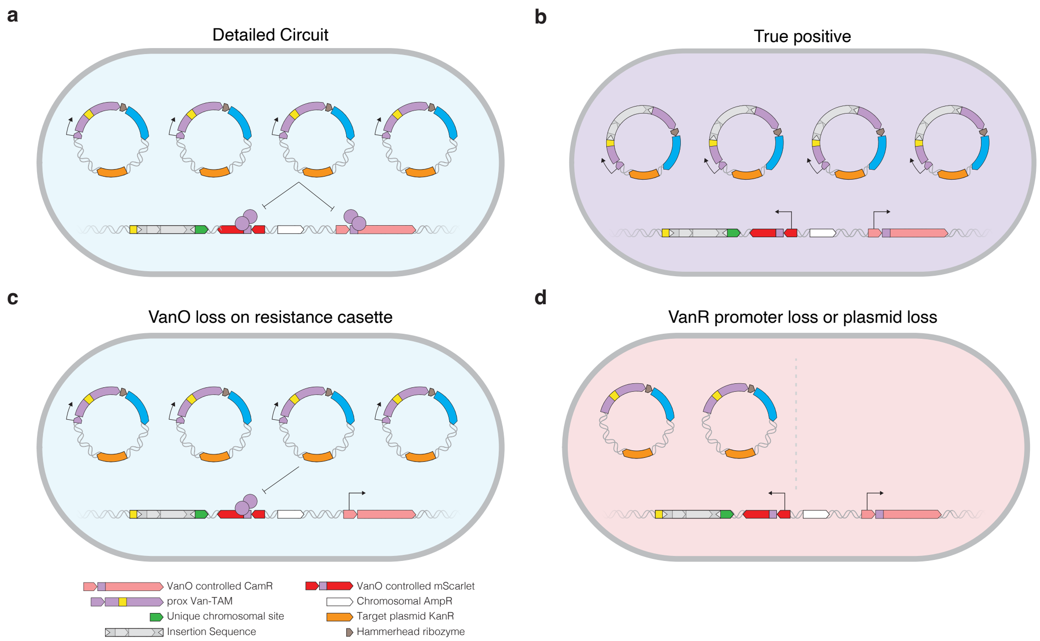


**Supplemental 4: Detailed circuit for transposon trap**

A) Transposon trap detailed circuit. Chloramphenicol resistance (purple) and mScarlet (red) are controlled by the vanillin operator (pink rectangle) that is contained in a polycistronic with mWatermelon (blue) linked with the hammerhead ribozyme (brown). Van-TAM is driven by the pVan promoter. The additional fluorophores enable control over some possible failure modes. B) Upon transposition into Van-TAM the VanR protein is disrupted allowing expression of mScarlet and chloramphenicol resistance. Surviving cells plated on chloramphenicol are only counted if they are also mScarlet and mWatermelon positive. C) Possible failure if the VanO controlling chloramphenicol resistance is lost or mutated. Surviving cells will only be mWatermelon positive. D) Possible failure if the promoter driving Van-TAM mutates (left) or if the plasmid is lost (right) and the cells become kanamycin resistant. Surviving cells will only be mScarlet positive. Any colonies detected that were single positive for mWatermelon or mScarlet were discounted in figure 3B.

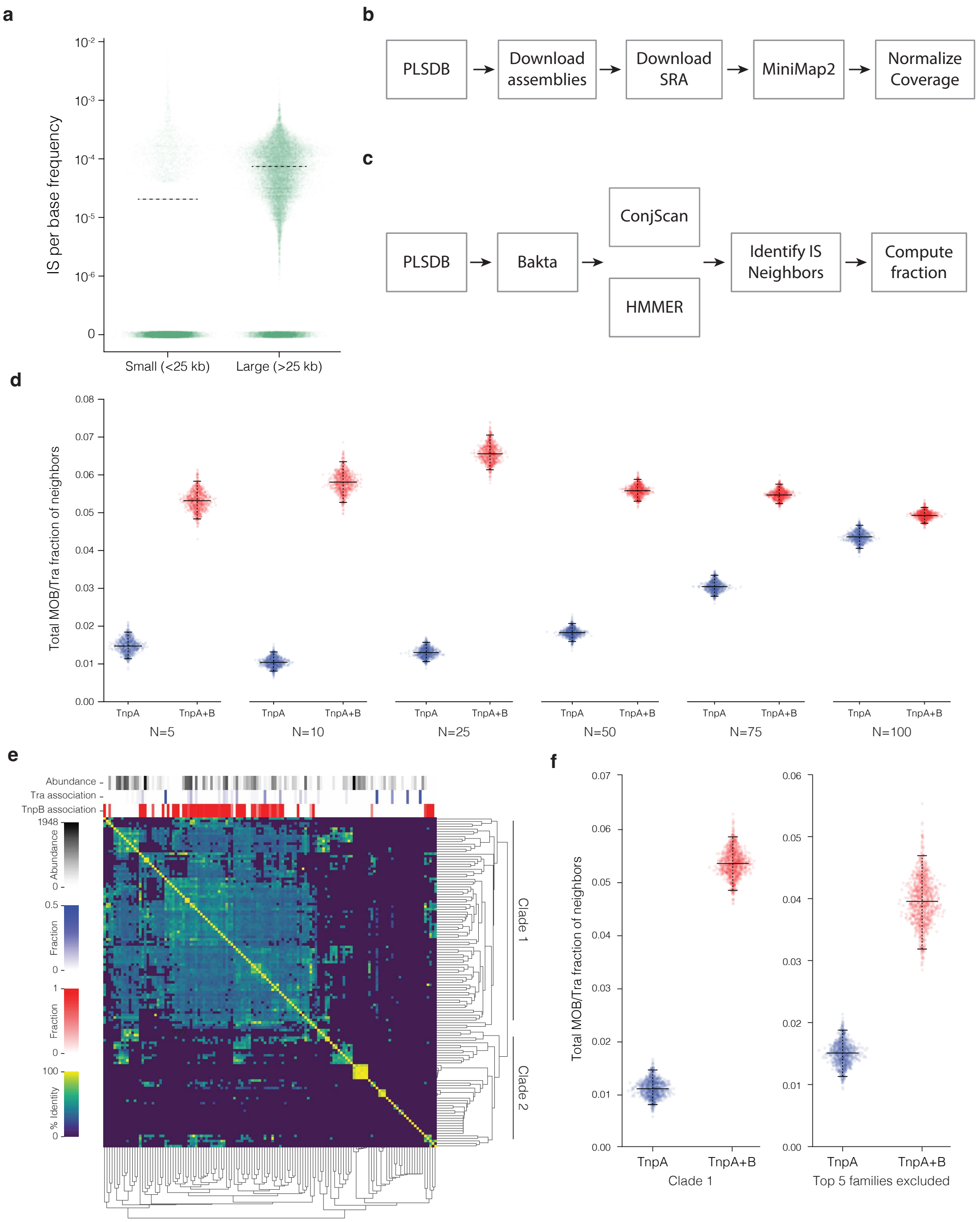


**Supplemental 5: Controlling for bias in neighborhood analysis**

A) IS per base frequency on small (<25 kb) and large (>25 kb) plasmids. The dotted line indicates the median. B) Pipeline to calculate coverage for plasmids with available short read sequencing in the PLSDB database. Sequence Read Archive files for all illumina sequenced assemblies in PLSDB were downloaded and aligned to the corresponding assembly with MiniMap2 to calculate relative coverage of the chromosome and plasmid C) Neighborhood analysis pipeline. Plasmids in PLSDB were homogeneously annotated with Bakta before identifying the mobilization genes with ConjScan and IS200 and IS605 elements with HMMER. The 5 flanking genes on each side (excluding the two most immediately adjacent genes to account for TnpB association in IS605 elements) of each IS were identified and the fraction of all that were ConjScan hits computed. D) Neighborhood analysis at different neighborhood windows. The difference between IS200 (blue) and IS605 (red) was significant until a window of 100 which would encompass nearly all the genes on every plasmid analyzed. E) Similarity clustering for neighborhood analysis. All TnpAs were extracted and clustered with MMSeqs2 at 50% AA similarity to determine broad families. Representatives from each family were pairwise aligned to create the similarity matrix and the relative abundance, plasmid association and TnpB association determined (top bars) for each family. TnpA families were clustered into two clades, one with greater TnpB association (predominantly IS605/IS607) and one without (IS200). F) Neighborhood analysis of IS200 (blue) and IS605 (red) when only analyzing TnpA from clade 1 (the TnpB associated clade) and excluding the top 5 most represented TnpA families.

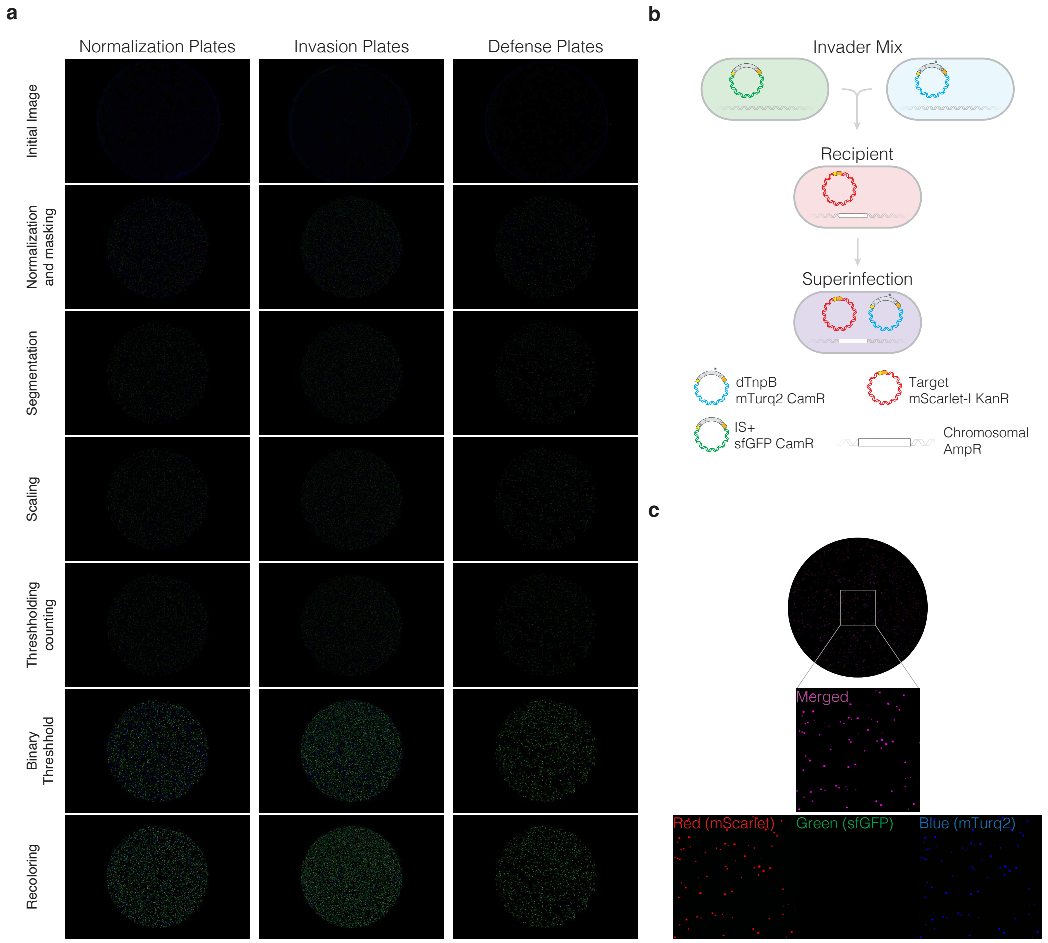


**Supplemental 6: Image analysis pipeline for conjugation invasion assays and F plasmid superinfection**

A) Image analysis pipeline. Initial images were normalized by the relative channel intensity, then masked to only include the interior of the plates. Colonies were identified with a watershed segmentation algorithm then scaled before thresholding applied to count for the analysis in figure 4E. For visualization a binary threshold was applied, and the channels maxed before recoloring. B) Superinfection of the F plasmid can be determined by selection on triple antibiotic plates. With a green tagged IS605 plasmid and a blue tagged IS605 harboring a dead TnpB, only the blue plasmid can coexist with the resident. Selected raw fluorescent images are depicted on the right enlarged at 3.5X.
